# Degradation of cytokinesis-specific Qa-SNARE KNOLLE is regulated by context-dependent ubiquitination

**DOI:** 10.64898/2026.05.13.724867

**Authors:** Misoon Park, Irina Droste-Borel, Boris Macek, Gerd Jürgens

## Abstract

In plant cytokinesis, the partitioning membrane is made by homotypic fusion of secretory vesicles, progressing in a centre-to-periphery direction. In Arabidopsis, this process is mediated by a cytokinesis-specific fusion machinery involving Qa-SNARE KNOLLE which is made during G2/M phase and degraded at the end of cytokinesis. Here we analyse how the turnover of KNOLLE protein is regulated. KNOLLE is ubiquitinated, which is best detected after combined treatment with inhibitors of endocytosis and de-ubiquitination. Site-directed mutagenesis of three clustered lysine residues prevented ubiquitination and internalisation, resulting in stable accumulation of KNOLLE at the plasma membrane in all cells of the seedling root. This is in stark contrast to the transient accumulation of wild-type KNOLLE in dividing cells only. Partial-substitution mutant lines revealed redundancy of lysine residues in both KNOLLE ubiquitination and turnover. KNOLLE ubiquitination resulted in K63-linked ubiquitin chains known to be involved in endocytosis whereas K48-linked chains were not detected. To explore the spatio-temporal conditions, we analysed KNOLLE ubiquitination in *cis*-SNARE and *trans*-SNARE complexes during membrane traffic and cell-plate formation. Our findings suggest that KNOLLE protein turnover is caused by a ubiquitination process that depends on successful membrane fusion generating the cell plate.

## Introduction

Protein turnover is a widespread mechanism of regulating the response to specific developmental or physiological situations such as hormone signaling, nutrient transport or pathogen defense (Hasegawa et al., 2024). A well-studied example is the transcriptional response to the plant signaling molecule auxin which is elicited by ubiquitination and subsequent proteasomal degradation of an IAA/AUX inhibitor that prevents an interacting ARF transcription factor from regulating the expression of target genes (de Roij et al., 2024). Another example is the boron-dependent ubiquitination and vacuolar degradation of the borate transporter BOR1 (Zhou et al., 2025; Yoshinari et al., 2021, 2024). Context-dependent protein turnover has also been detected in the cytokinesis of the flowering plant Arabidopsis. Specifically, the Qa-SNARE KNOLLE accumulates in a cytokinesis-specific manner and is required for membrane fusion of secretory vesicles that form the partitioning membrane known as cell plate (Lukowitz et al., 1996; Lauber et al., 1997). KNOLLE forms *cis*-SNARE complexes with promiscuous SNARE partners at the endoplasmic reticulum (ER). These complexes are delivered via the Golgi stack to the *trans*-Golgi network (TGN) where they are sorted into forming vesicles destined for the cell division plane (Reichardt et al., 2007; Richter et al., 2014; Karnahl and Park et al., 2017). Following disassembly of the *cis*-SNARE complexes through the action of the NSF ATPase and its adaptor αSNAP2, the Sec1/Munc18 (SM) protein KEULE assists in the formation of *trans*-SNARE complexes that mediate fusion between neighbouring vesicles or between later-arriving vesicles and the margin of the laterally expanding cell plate (Park et al., 2012; Park et al., 2023). At the end of cytokinesis, KNOLLE protein is endocytosed and targeted to the vacuole for degradation (Reichardt et al., 2007). In an Arabidopsis mutant expressing a truncated subunit of the endocytosis-specific adaptor complex TPLATE, KNOLLE resided longer at the cell plate than in wild-type cytokinesis and its ubiquitination was detected by immunoblotting (Grones et al., 2022).

Here, we show that KNOLLE needs to be ubiquitinated for being endocytosed and targeted to the vacuole for degradation. We identify 3 lysine residues that contribute to KNOLLE ubiquitination required for rapid turnover at the end of cytokinesis. We also demonstrate that ubiquitination of KNOLLE involves the formation of K63-linked ubiquitin chains which serve as a signal for endocytosis and degradation in the vacuole. In addition, KNOLLE appears not to be ubiquitinated when trapped in SNARE complexes, which suggests that membrane fusion generating the cell plate is required for ubiquitination-dependent degradation of KNOLLE.

## Results

### Detection of ubiquitinated forms of KNOLLE

Presuming that KNOLLE turnover was ubiquitin-dependent, we employed anti-ubiquitin antibody to detect ubiquitinated GFP:KNOLLE fusion protein in seedling extracts, following immunoprecipitation with the GFP-trap beads (Figure 1). For comparison, we analysed extracts from wild-type and NSF:GFP transgenic seedlings. The KNOLLE-enriched precipitate showed a distinct pattern of anti-ubiquitin-detectable bands that was not present in the other samples (bracket, Figure 1, *bottom*). There were several weak bands presumably representing ubiquitinated forms of KNOLLE. We reasoned that ubiquitination might result in fast endocytosis and targeting to the vacuole for degradation, which would explain the low level of ubiquitinated KNOLLE accumulation. In addition, this diminishing effect might be enhanced by the action of de-ubiquitinating enzymes (reviewed by Vogel and Isono, 2024a,b). In order to define the bands more clearly, we attempted to increase the amount of ubiquitinated KNOLLE with specific treatments (Figure S1). We treated *GFP:KNOLLE* transgenic seedlings with wortmannin, which we had shown to impair endocytosis (Reichardt et a., 2007). Compared to the mock control, the treated samples gave slightly enhanced signals (Figure S1A). N-ethylmaleimide (NEM) treatment affects the activity of general de-ubiquitinating enzymes (Loch et al., 2011; Chen et al., 2018). Its effect on detectable bands of ubiquitinated KNOLLE was a little stronger than the effect of wortmannin. By contrast, the combined treatment of seedlings with both wortmannin and NEM, together with the presence of NEM during protein extraction and immunoprecipitation, caused approximately 15-fold enhancement such that strong signals were detected with both, anti-ubiquitin and anti-KNOLLE antibodies (Figure S1B; double arrows).

**Figure 1.**
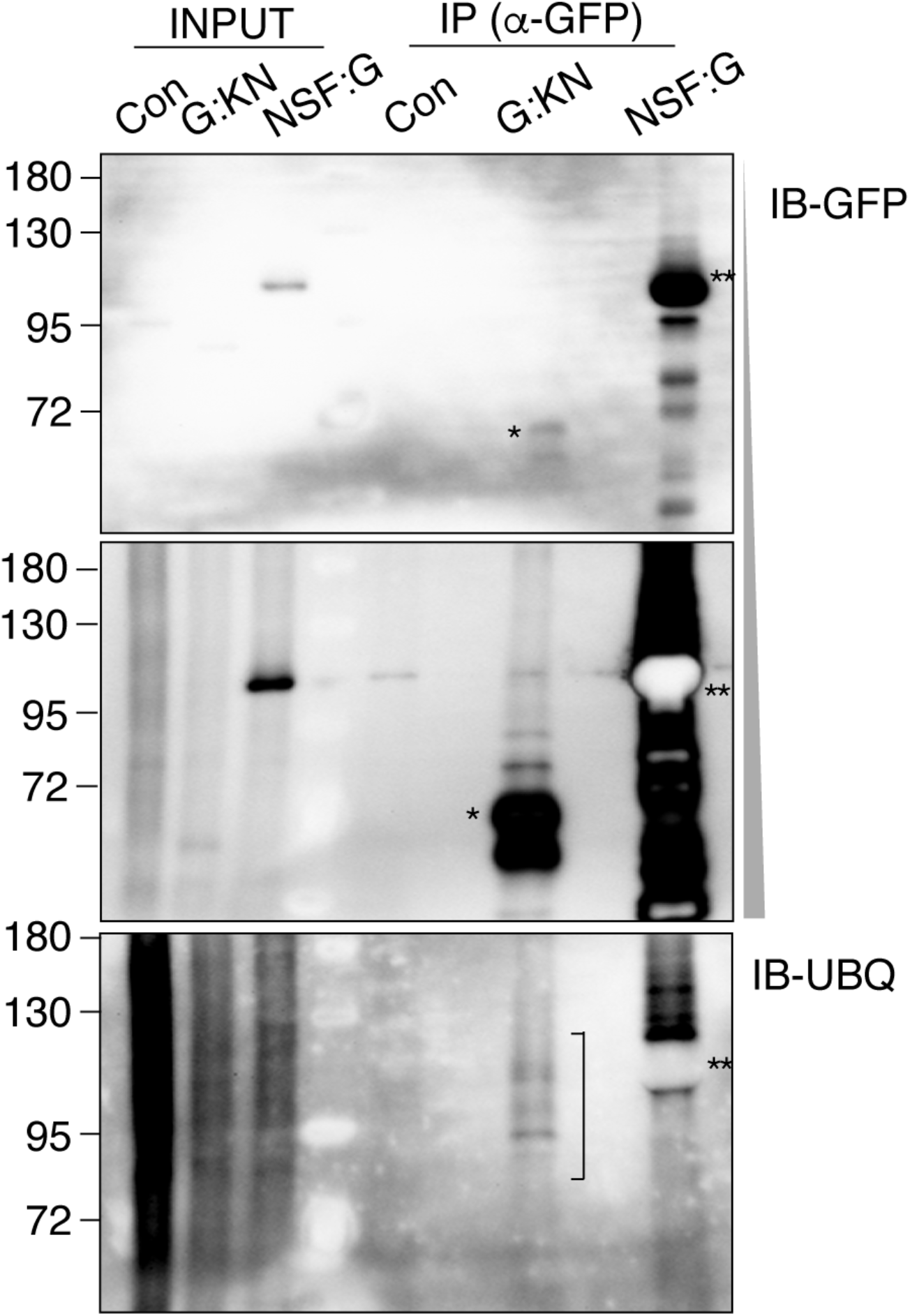
Detection of ubiquitinated KNOLLE protein. Protein extracts from non-transformed wild type (Con), GFP-tagged KNOLLE (G:KN) or GFP-tagged NSF (NSF:G) transgenic seedlings were immunoprecipitated with GFP-trap beads (IP (α-GFP)) and the precipitated proteins separated by SDS-PAGE and probed for GFP (IB-GFP: shorter exposure, *top;* longer exposure, *middle*) and for ubiquitin (IB-UBQ, *bottom*). INPUT, total protein extract. Note strong accumulation of GFP:KNOLLE below 72 kDa (G:KN, asterisk, *top* and *middle*) and NSF:GFP (NSF:G near 120 kDa, double asterisks, *top* and *middle*) and 4 or 5 weak bands of ubiquitinated G:KN between 72 and 130 kDa that were absent from the NSF:G sample (bracket; *bottom*).

Having established that KNOLLE is indeed ubiquitinated, we next addressed the biological significance of this modification. To that end, we sought to identify lysine residues that might be subject to covalent linkage with ubiquitin. Immunoprecipitation followed by mass spectrometry (IP-MS) identified a single ubiquitinated lysine residue K_223_ in KNOLLE protein (Figure S2A). We then engineered a substitution variant of KNOLLE named KN^K1R^ in which that lysine residue (K_223_) was replaced by arginine (R) (Figure S2B) and generated transgenic Arabidopsis expressing the venusYFP (YFP)-tagged variant from a *KNOLLE* expression cassette (Figures S2C-F and S3; Müller et al., 2003). YFP-tagged KN^K1R^ showed a wild-type pattern of fluorescence in the seedling root. Thus, K_223_ seems not to be involved in the ubiquitination process that is required for KNOLLE turnover.

### Lysine residues involved in ubiquitination and turnover of KNOLLE

Because no other ubiquitinated residue had been detected by IP-MS we took a different approach, inspired by a study of human syntaxin 3 (Stx3). That protein has 6 lysine residues in its juxtamembrane region and appears to play a ubiquitination-dependent role in trafficking to apical exosomes in epithelial cells (Giovannone et al., 2017). By aligning the sequences of members of the Arabidopsis SYP1 family of plasma membrane-localised Qa-SNAREs (aka syntaxins) with Stx3, we identified 3 potential ubiquitination residues near the C-terminal membrane anchor of KNOLLE (Figure S2B). We then generated transgenic substitution lines expressing different YFP-tagged KNOLLE variants. All 3 lysine residues K_272_, K_275_ and K_283_ were substituted by arginine in Y:KN^K234R^, only residue K_272_ in Y:KN^K2R^ or the complementary 2 residues K_275_ and K_283_ in Y:KN^K34R^ were replaced by arginine (Figure 2A; Figure S3).

**Figure 2.**
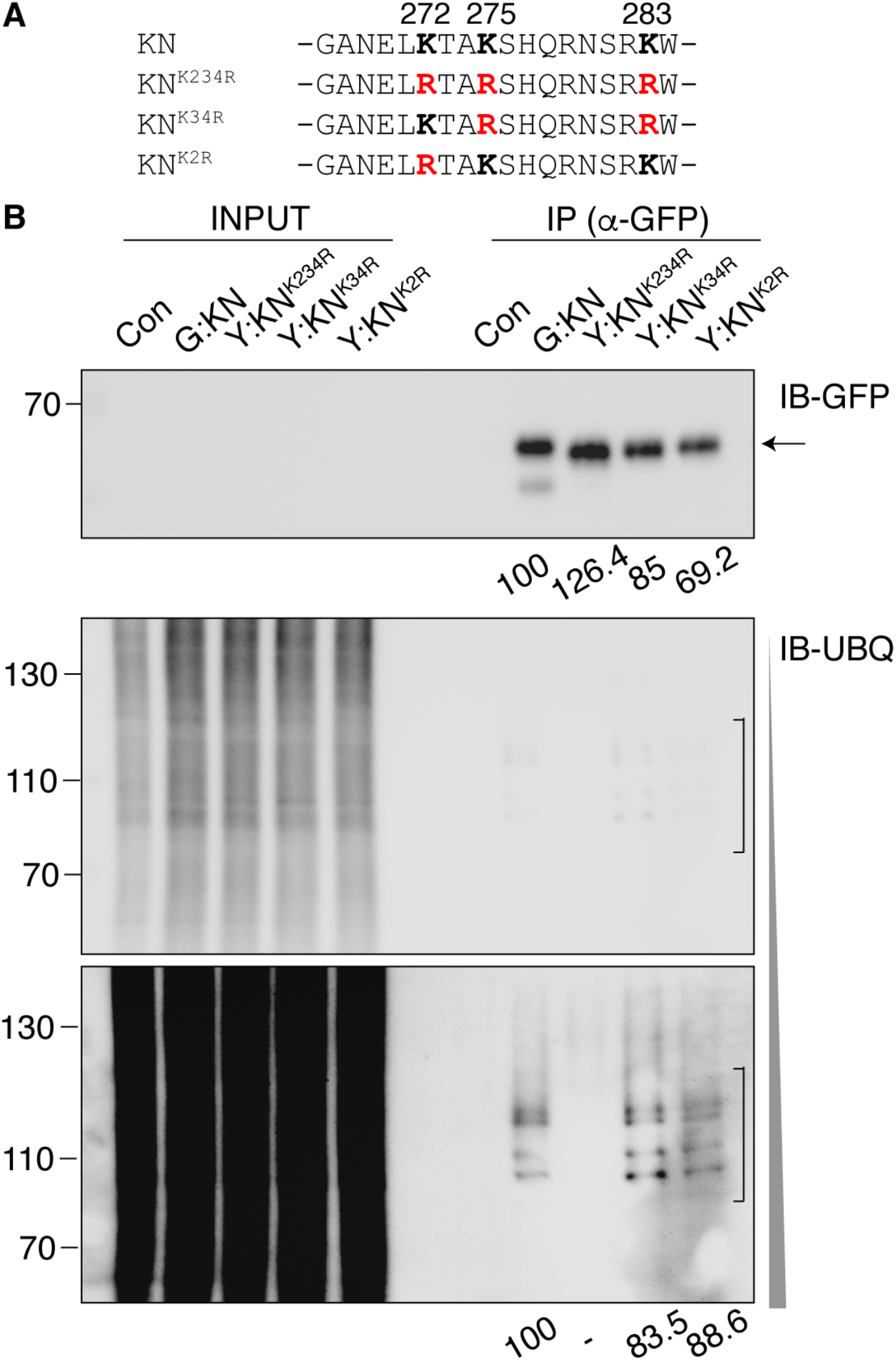
Effect of lysine-to-arginine substitutions on KNOLLE ubiquitination. (**A**)Juxtamembrane segment of KNOLLE and its substitution variants. Substituting arginine (R) residues are highlighted in red; positions of relevant amino acid residues are indicated at the top. See also Figure S2B for sequence alignment of KNOLLE homologs in Arabidopsis. (**B**)Immunoprecipitated wild-type GFP-tagged KNOLLE (G:KN) and its YFP-tagged substitution variants (Y:KN^K234R^, Y:KN^K34R^, Y:KN^K2R^) as well as non-transformed wild-type (Con) were probed for ubiquitinated forms by immunoblotting with anti-ubiquitin antibody (IB-UBQ). INPUT, total protein extract; IP (α-GFP), immunoprecipitation with GFP-trap beads. Molecular size markers in kDa (left). Note that several ubiquitinated bands (brackets) are completely absent in Y:KN^K234R^ variant. Arrow, proteins of GFP:KNOLLE and its variants (*top* and *bottom*). Numbers below top panel and prolonged-exposure panel indicate signal intensities relative to wild-type GFP-tagged KNOLLE (G:KN).

Any effects that the K-to-R substitutions might have on the ubiquitination of KNOLLE were assessed by probing immunoprecipitated KNOLLE variants with ubiquitin-specific antibodies (Figure 2B). In contrast to the ubiquitin-positive KNOLLE wild-type protein, the triple-substitution variant KN^K234R^ lacked detectable ubiquitin modification completely. However, each of the other two variants still displayed a seemingly identical pattern of ubiquitination bands to that of wild-type KNOLLE. These results indicate that no other lysine residues than those 3 clustered in the juxtamembrane region contribute to detectable KNOLLE ubiquitination. In addition, at least 2 of the 3 relevant lysine residues can be ubiquitinated. Furthermore, the levels of ubiquitination appear to be comparable between the partial-substitution variants and wild-type GFP-tagged KNOLLE as the semi-quantitative assessment showed no obvious differences (Figure 2B). This suggests that a single lysine residue within the cluster might be sufficient for the full ubiquitination level of KNOLLE protein.

The biological significance of KNOLLE ubiquitination was assessed by confocal imaging of seedling roots expressing the different KNOLLE substitution variants (Figure 3A-L). With all 3 lysine residues substituted (KN^K234R^), KNOLLE labelled the plasma membrane of all root cells, thus accumulating like the plasma membrane marker, SynaptoRed^™^ C2 (Figure 3D-F). By contrast, the partial-substitution variants of KNOLLE, Y:KN^K34R^ and Y:KN^K2R^, essentially showed the wild-type pattern of accumulation in dividing cells (compare Figure 3 panels G-I and J-L with A-C). Measurement of the relative signal intensity at the plasma membrane versus the cell-division plane clearly showed that Y:KN^K234R^ was more stable than the two partial-substitution variants and wild type KNOLLE (Figure 3Q). These results indicate that the remaining lysine residues, K_272_ in Y:KN^K34R^ and K_275_ and/or K_283_in Y:KN^K2R,^, can still mediate KNOLLE turnover. Although high-level expression of Y:KN^K2R^ appeared to delay turnover of KNOLLE (Figure S4M-O), it did not completely block its turnover, unlike Y:KN^K234R^. This result supports our view that KNOLLE turnover is primarily governed by its ubiquitination, although the dynamics of turnover might be modulated by the level of protein accumulation.

**Figure 3.**
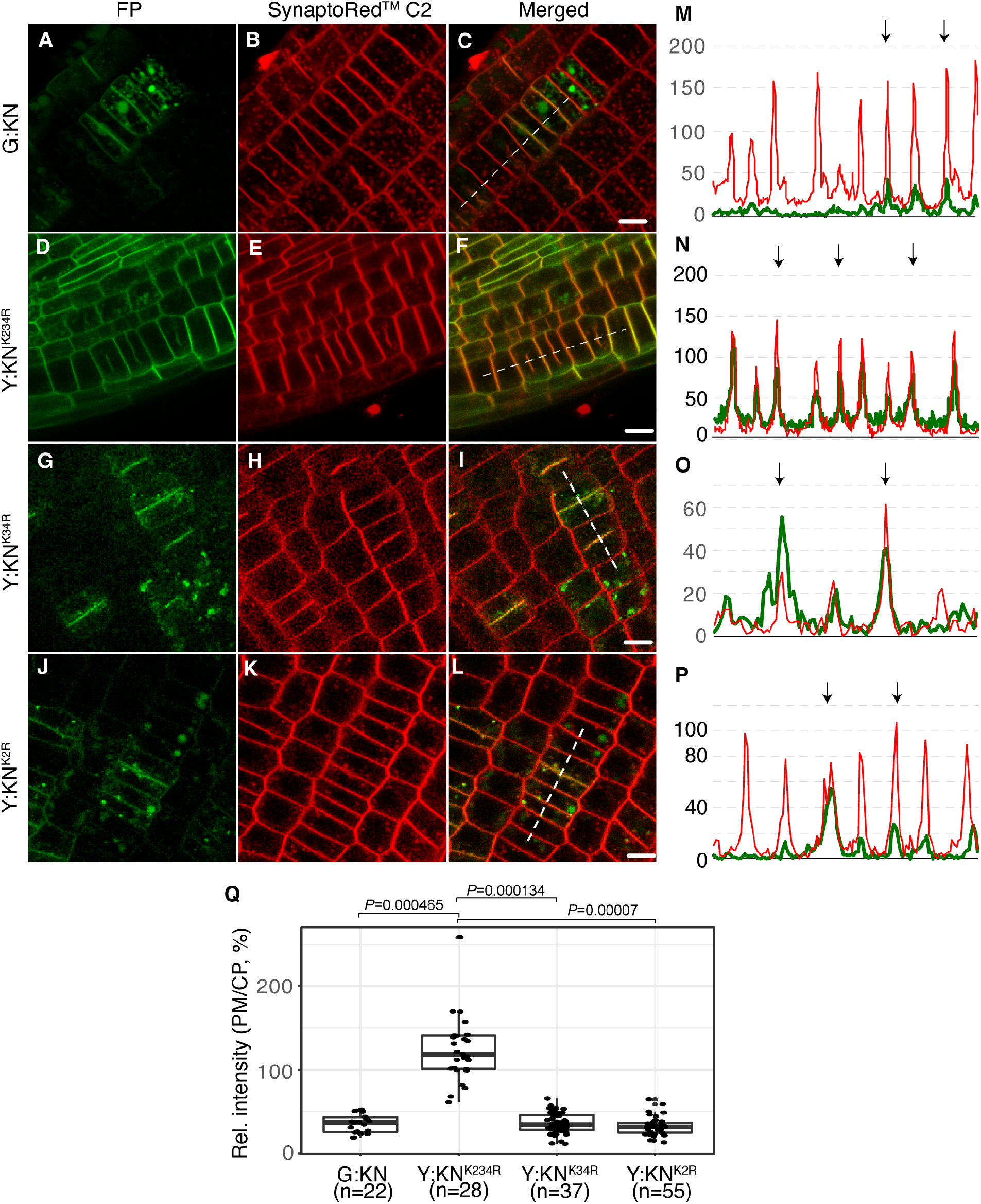
Subcellular accumulation of KNOLLE protein with K-to-R substitutions. (**A-L**) Live imaging of tagged KNOLLE variants in seedling root cells: (**A-C**) GFP:KNOLLE; (**D-F**), YFP: KN^K234R^; (**G-I**), YFP:KN^K34R^; (**J-L**), YFP:KN^K2R^. Scale bars, 5 μm (**C, F, I, L)**. (**M-P**) Line scans of KNOLLE variants (green) and plasma membrane marker SynaptoRed^™^C2 (red) along the dashed lines marked in (**C, F, I, L**). Arrows at the top indicate cell plates. Note clear difference between the fully substituted variant Y:KN^K234R^ and all the other variants (– tagged KNOLLE might in general take somewhat longer to be internalised than endogenous untagged KNOLLE protein). (**Q**) Relative signal intensity of tagged KNOLLE variants in dividing cells expressed as ratio of cell-plate signal to plasma-membrane signal (PM/CP, in %). n, Number of the counted cells. Statistical analysis with unpaired Student’s t-test and *P* values with two-tailed tests.

Upon seedling treatment with the fungal toxin brefeldin A (BFA), KNOLLE accumulates in endosomal BFA compartments, which can occur both on the way from the TGN to the division plane and after endocytosis from the cell plate on the way to the vacuole (Reichardt et al., 2007; Richter et al., 2014). All substitution variants of KNOLLE accumulated in the BFA compartments in dividing cells, implying that the ubiquitination potential does not influence KNOLLE traffic from the TGN to the division plane (Figure S5). No obvious accumulation of Y:KN^K234R^ in the BFA compartments was observed in interphase cells while the protein was present at the plasma membrane, which supports our view that KNOLLE is not endocytosed in the absence of ubiquitination.

It should be noted that stabilised KNOLLE protein is active and can fully rescue *knolle*^*X37-2*^ deletion mutant plants (Table S1). This rescue also implies that KNOLLE being present on the plasma membrane in post-mitotic cells does not interfere with any essential process (see Discussion).

### KNOLLE ubiquitination involves K63-linked ubiquitin chains

The size shift of ubiquitinated KNOLLE by up to approximately 60 kDa would be compatible with ubiquitin chains being covalently linked to one or more of the 3 lysine residues in the juxtamembrane region (see Figure 1). Endocytosis of plasma membrane proteins has been reported to involve K63-linked ubiquitin chains (Saeed el., 2023; Hasegawa et al., 2024). We used K63 and K48 linkage-specific antibodies to assess the presence of ubiquitin chains on KNOLLE protein and to distinguish between two major types of chains. In this experiment, we also compared mock-treated samples with samples from seedlings treated with wortmannin and NEM (Figure 4). The anti-K48 antibody did not detect ubiquitin in the immunoprecipitated fraction of GFP:KNOLLE. By contrast, the K63 antibody detected a broad band of approximately 120 kDa which was enhanced in the treated sample compared to the mock-treated one (Figure 4, lower panel). No such band was detected in the samples from the triple-substitution line YFP:KN^K234R^. These results indicate that ubiquitinated KNOLLE bears at least one K63-linked ubiquitin chain.

**Figure 4.**
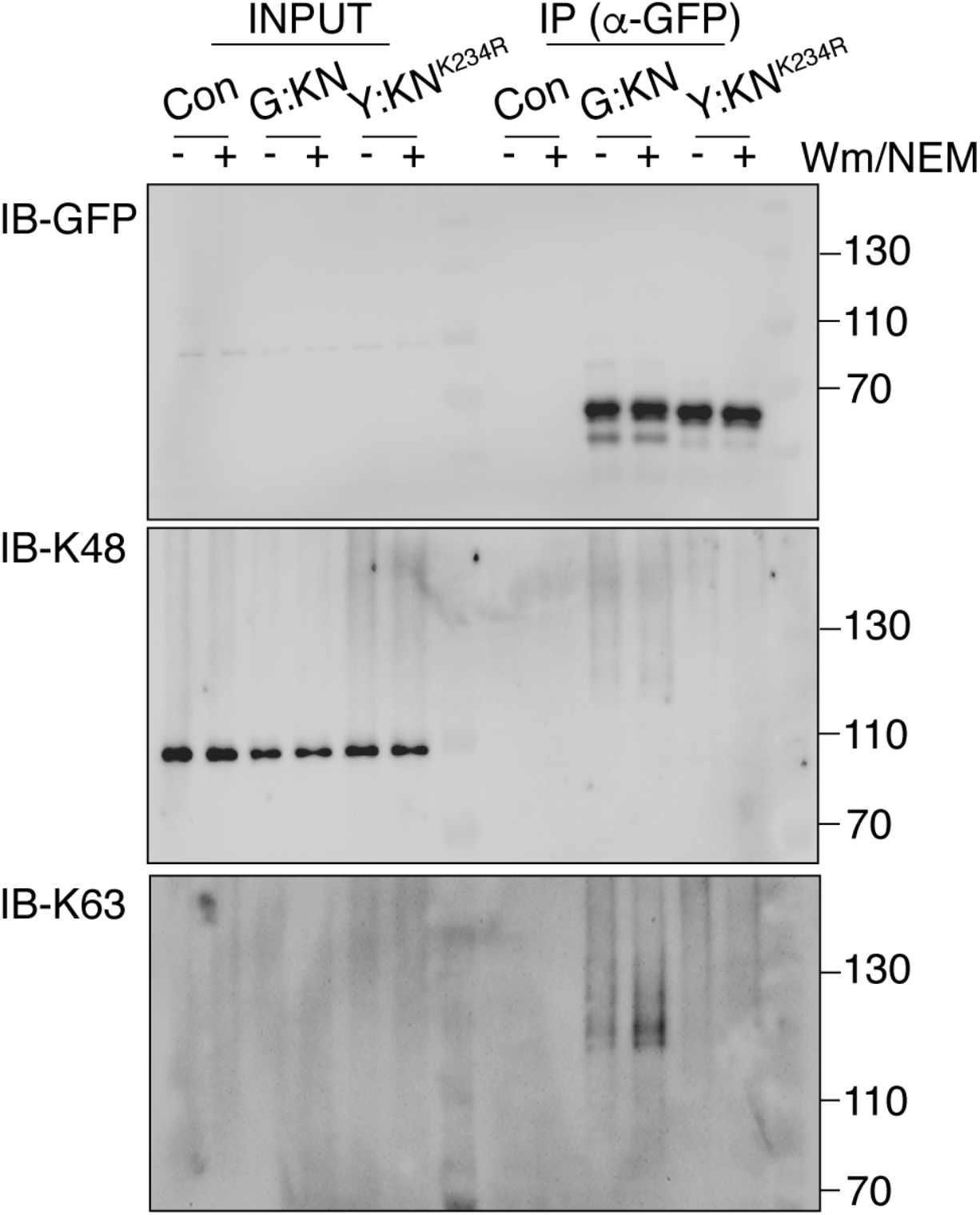
Detection of specific ubiquitin-chain linkage. Protein extracts from seedlings treated with wortmannin and NEM were immunoprecipitated with the GFP trap (IP (α-GFP)) and the enriched fractions probed with linkage-specific antibodies as indicated on the right. INPUT, total protein extract; -, mock-treated seedlings. Con, non-transgenic control; G:KN, GFP-tagged KNOLLE wild-type; Y:KN^K234R^, YFP-tagged triple-substitution variant of KNOLLE; Numbers, protein sizes in kDa (*right*).

### Context-dependent ubiquitination of KNOLLE

Having established the functional significance of ubiquitination in KNOLLE turnover, we addressed when and where KNOLLE ubiquitination might occur. The experiment was to provide a reference by preventing the formation of cytokinetic membrane vesicles that normally deliver KNOLLE from the TGN to the plane of cell division. This trafficking step can be inhibited by BFA treatment in *big3* mutant seedlings in which the late secretory pathway mediated by redundant ARF-GEFs BIG1 to BIG4 is rendered BFA-sensitive (Richter et al., 2014; Karnahl and Park et al., 2017). In the *big3* mutant seedlings, we expressed the GFP-tagged KNOLLE partner SNAP33 in an estradiol-inducible fashion. Seedlings expressing the GFP:KNOLLE fusion were analysed as control samples in a parallel set of experiments. All protein extracts were immunoprecipitated with GFP-trap beads and then probed with anti-GFP, anti-KNOLLE and anti-ubiquitin antibodies (Figure 5).

**Figure 5.**
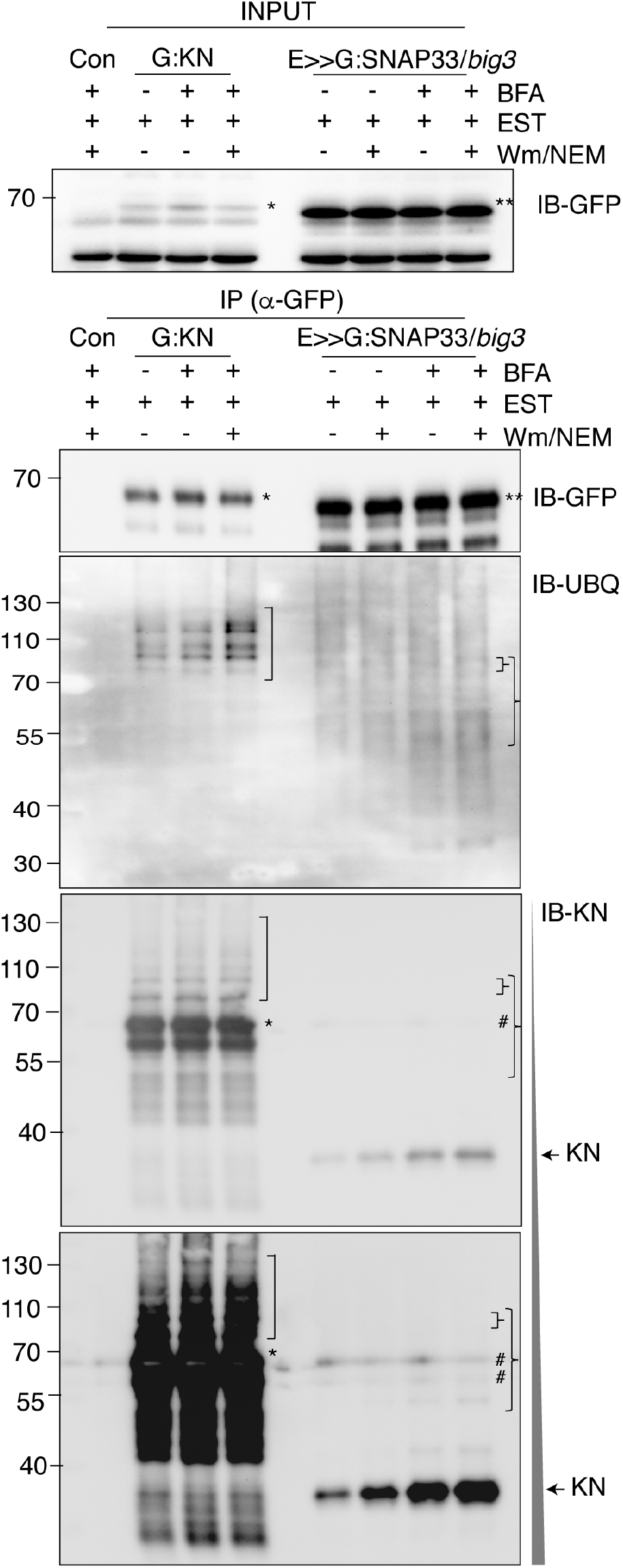
Inhibition of membrane traffic and its effect on KNOLLE ubiquitination. KNOLLE as part of SNARE complex (E>>G:SNAP33/*big3*, estradiol-induced, +EST) either trapped at TGN (+ brefeldin A, BFA) or trafficked to the cell division plane (-BFA) was tested for ubiquitination. Lines for comparison: Con, non-transgenic control; G:KN, GFP-tagged KNOLLE.Protein extracts from seedlings treated with wortmannin (Wm) and NEM (+Wm/NEM) or untreated (-Wm/NEM) were immunoprecipitated with the GFP trap (IP(α-GFP)) and the enriched fractions probed with anti-GFP, anti-ubiquitin and anti-KNOLLE antibodies; bottom panel, prolonged exposure. Expected sizes: GFP:KNOLLE, 65 kDa; (UBQ)n-GFP:KNOLLE, 80-120 kDa (UBQ moiety 15-60 kDa); endogenous KNOLLE, 33 kDa; (UBQ)n-KNOLLE, 50-100 kDa. Brackets, ubiquitinated bands of GFP:KNOLLE; wide braces, range of (UBQ)n of KNOLLE between 50 and 100 kDa; narrow braces, the highest (UBQ)n band of KNOLLE at approximately 100 kDa. Asterisk, GFP:KNOLLE; double asterisks, GFP:SNAP33; hashtags, cross-reacting bands of anti-KNOLLE antiserum; numbers on the left, protein sizes in kDa.

The GFP:KNOLLE samples yielded the expected results. KNOLLE ubiquitination was detected and the signals were enhanced by the combined treatment with wortmannin and NEM, regardless of BFA treatment (because of BFA-resistant ARF-GEF BIG3) (compare with Figure S1). By contrast, the GFP:SNAP33 immunoprecipitates from BFA-treated seedlings did not yield ubiquitinated KNOLLE, although a comparable amount of KNOLLE was detected (bottom panel in *Figure 5*). This observation was confirmed by treatment of the GFP:SNAP33 transgenic seedlings with wortmannin and NEM which did not give any indication of KNOLLE ubiquitination either. These results suggest that KNOLLE being part of the *cis*-SNARE complex is not ubiquitinated. Interestingly, protein extracts from GFP:SNAP33 transgenic seedlings not treated with BFA also did not display ubiquitinated KNOLLE in their GFP:SNAP33 immunoprecipitates (Figure 5). In this situation, trafficking to the plane of cell division results in NSF/αSNAP2-driven disassembly of *cis*-SNARE complexes and assembly of KEULE-assisted *trans*-SNARE complexes required for cell-plate formation. The absence of detectable ubiquitinated KNOLLE thus seems to suggest that KNOLLE complexed with SNARE partners in general might not be ubiquitinated (see Discussion).

## Discussion

Plant cytokinesis employs a specific membrane-fusion machinery which lays down the partitioning membrane in a centrifugal fashion. The key factor Qa-SNARE KNOLLE is only made during G2/M phase and turned over at the end of cytokinesis (Lauber et al., 1997; Reichardt et al., 2007). The regulation of KNOLLE synthesis is fairly well understood, involving MSA elements and R1R2R3-Myb transcription factors (Haga et al., 2007). By contrast, much less is known about the turnover of KNOLLE at the end of cytokinesis, although endocytosis and targeting to the vacuole for degradation have been described (Reichardt et al., 2007; Boutté et al., 2010). Slowed-down endocytosis caused by reduced activity of the TPLATE complex enabled an ubiquitinated form of KNOLLE to be detected (Grones et al., 2022).

### Lysine redundancy in KNOLLE ubiquitination and turnover

Our results indicate that lysine residues in the juxtamembrane region, i.e. within a stretch of 15 amino acid residues near the membrane anchor, are essential to KNOLLE ubiquitination that is required for endocytosis and degradation in the vacuole. One or more of these lysine residues appear to have covalently linked ubiquitin chains of the K63-type, which seems to be a general signal for endocytosis (Saeed et al., 2023). Of the three lysine residues, at least two can be ubiquitinated. Each ubiquitination appears to be sufficient to mediate KNOLLE turnover of the partial-substitution variants. Moreover, there was no obvious difference in the pattern and intensities of ubiquitinated KNOLLE bands between the wild-type and the partial-substitution variants. These observations suggest redundancy of the clustered lysine residues in KNOLLE ubiquitination and turnover.

Redundancy would also be consistent with an evolutionary survey of KNOLLE sequences. KNOLLE only exists in angiosperms and seems to have originated in the context of double fertilisation which produces a cellularising endosperm in addition to a zygotic embryo (Park et al., 2018). Of 182 angiosperm species representing basal angiosperms, monocots, magnoliids and eudicots, 97 make KNOLLE proteins with all 3 K residues, 79 with 2 K residues, and only 6 with 1 K residue in the juxtamembrane region. Thus, there is clear selection on the lysine residues to be ubiquitinated. Nonetheless, non-degradable KNOLLE (KN^K234R^) is active and has no deleterious effects on development or reproduction. Why then is KNOLLE normally turned over at the end of cytokinesis? Tentative explanations might be: (1) Nitrogen supply is not limited under laboratory conditions so the mutant plants can afford not to recycle amino acids from stable KNOLLE protein. The situation might be very different in the wild. (2) Although the stable variant of KNOLLE appears not to be deleterious, other potential substrates of a cytokinesis-specific protein degradation mechanism might interfere with essential processes if still present in post-mitotic cells.

### Context-dependent degradation of KNOLLE

Newly-synthesised KNOLLE protein and its SNARE partners are assembled into a *cis*- SNARE complex at the ER membrane and stay in that complex until they have been delivered to the plane of cell division (Karnahl and Park et al., 2017). Here, NSF ATPase and its αSNAP2 adapter disassemble the *cis*-SNARE complex (Park et al., 2023). SNARE complex disassembly would normally result in monomeric Qa-SNARE as a consequence of its N-terminally located alpha-helices folding back onto the SNARE domain, which prevents re-formation of the SNARE complex (Yoon and Munson, 2018). However, SM protein KEULE captures Qa-SNARE KNOLLE by its linker, promoting the formation of *trans*-SNARE complex, which is required for membrane fusion during cell-plate formation (Park et al., 2012). Thus, KNOLLE essentially remains in SNARE complexes from its synthesis at the ER up to the formation of the cell plate during cytokinesis. The implication of this scenario is that the ubiquitination machinery may only be able to access its substrate after KNOLLE has performed its role in cell plate formation. It is conceivable that the SNARE complex sterically hinders the ubiquitination machinery from accessing its target KNOLLE. The conclusion that KNOLLE ubiquitination is constrained in place and time is also consistent with our earlier finding that KNOLLE expression from the constitutively active *35S* promoter results in stable accumulation at the plasma membrane of non-dividing root hair cells (Völker et al., 2001).

Taken together, our results suggest a model of how the turnover of KNOLLE might be coordinated with the progression of cytokinesis (Figure 6). In essence, following the disassembly of *cis*-SNARE complexes by NSF and αSNAP2, SM protein KEULE mediates the transition to *trans*-SNARE complexes which promote membrane fusion during cell-plate formation. It is in this context that KEULE interacts with KNOLLE and thus plays an essential role in membrane fusion (Park et al., 2012). In other words, KEULE plays a role in cytokinesis before membrane fusion but not thereafter. In view of the available evidence, we would like to propose that KNOLLE-mediated cell-plate formation is an essential step for timely KNOLLE degradation. Membrane fusion might activate relevant ubiquitin ligase(s) that can capture monomeric KNOLLE on the cell plate after disassembly of the post-fusion *cis*-SNARE complex in the absence of KEULE (Figure 6). Once ubiquitinated, KNOLLE is endocytosed and targeted to the vacuole for degradation.

**Figure 6.**
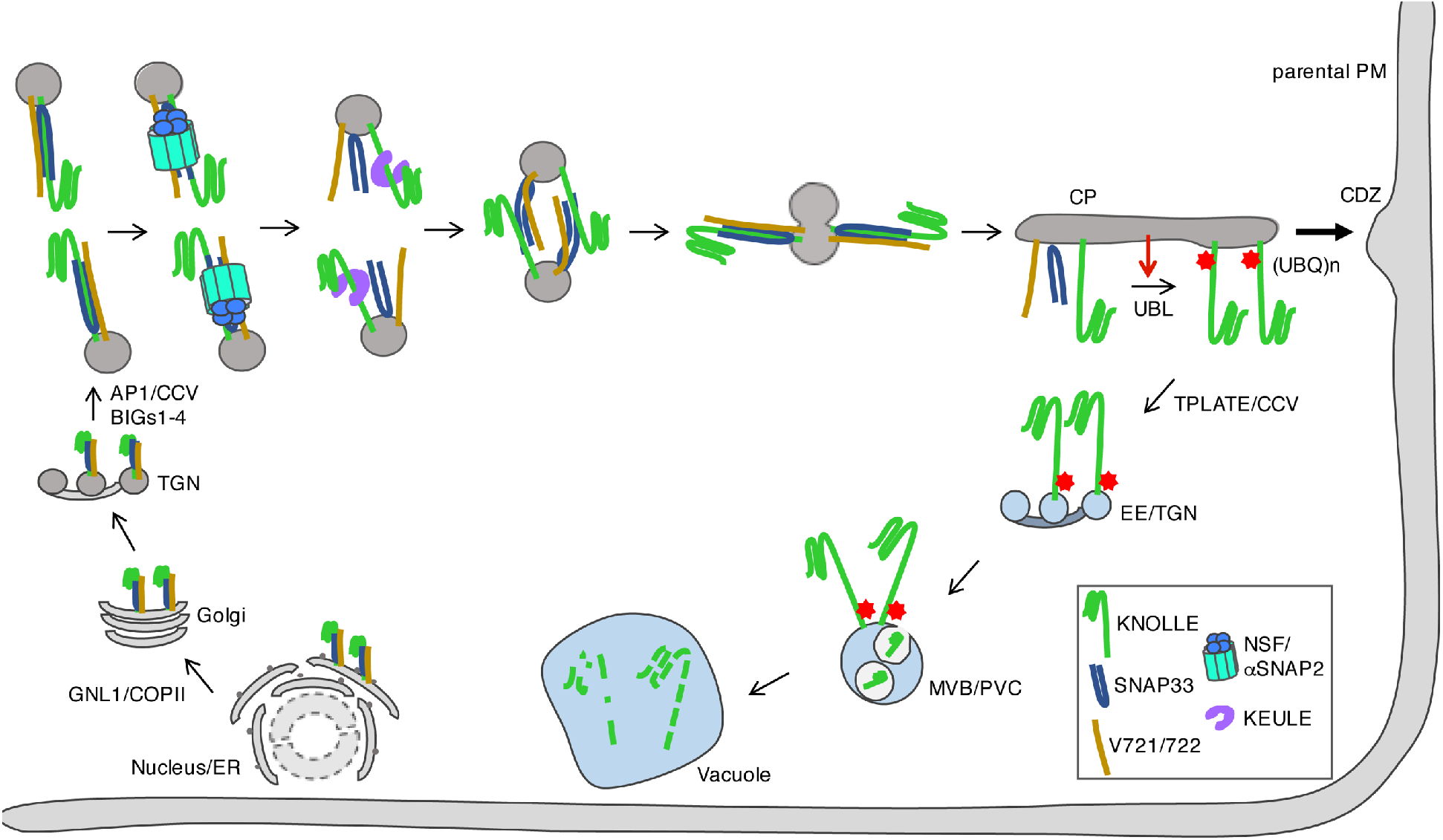
KNOLLE ubiquitination in the spatio-temporal context of cytokinesis (model) Disassembly of *cis*-SNARE complexes (KNOLLE, green; SNAP33, dark blue; VAMP721/722 (V721/722), brown) on TGN-derived membrane vesicles by NSF ATPase (turquiose) and αSNAP2 (blue) adapter results in *cis*-*trans* conversion of SNARE complexes in the presence of SM protein KEULE (purple), which promotes membrane fusion. As a consequence, the forming cell plate harbours *cis*-SNARE complexes which are again disassembled by NSF and α-SNAP. In the absence of KEULE and possibly induced by a signal from the cell plate (red arrow), a ubiquitin ligase (UBL) covalently links ubiquitin with 3 lysine residues of KNOLLE (red stars) which is thus marked for rapid endocytosis and targeting to the vacuole via EE/TGN (early endosomes or TGN) and MVB/PVC (multivesicular bodies/prevacuolar compartments) for degradation.

## Experimental procedures

### Plant lines and growth conditions

*Arabidopsis thaliana* was grown on soil at 23°C or 18°C in a long-day condition (16hr/8hr). Wild-type Columbia (Col-O) or heterozygous plants of *knolle*^*X37-2*^ were transformed with *Agrobacterium tumefaciens*, using the floral-dip method (Clough and Bent, 1998). T1 seedlings were selected by spraying three times 1:1000 diluted BASTA (183 g/l glufosinate; AgrEvo, Düsseldorf, Germany) or PPT (phosphinothricin, 15 ug/ml, Sigma-aldrich). Selection was confirmed by PCR genotyping with forward primer vYFP640for and gene-specific reverse primers (Table S2). *pKNOLLE::GFP:KNOLLE* (El Kasmi and Krause et al., 2013), *pMDC7::NSF:GFP* (Park et al., 2023) and *pMDC7::GFP:SNAP33* in the *big3* homozygous background (Karnahl and Park et al., 2017) were also used in this study.

### Complementation tests

T2 seeds were germinated on solid medium (1/2 MS, 0.1% MES, 1% sucrose, pH5.6) in the same growth condition as described above. Segregation of antibiotics resistance was counted on PPT (15 µg/ml)-supplemented medium. Rescue of *knolle*^*X37-2*^ mutant was determined by counting the percentage of *knolle*^*X37-2*^ in the segregating T2 population and with PCR using primers CIIIext and DIIIext for *knolle*^*X37-2*^, and DIIIext and DEL for *KNOLLE* (Table S2).

### Molecular cloning

For *pGIIB-KN::venusYFP:KN*^*1R*^, an upstream fragment was PCR amplified with primers MP67 and KN_K1R_rev and a downstream fragment with primers KN_K1R_for and MP68. For *pGIIB-KN::venusYFP:KN*^*34R*^, an upstream fragment was PCR amplified with primers MP67 and KN_K3R_rev and a downstream fragment with primers KN_K4R_for and MP68. For *pGIIB-KN::venusYFP:KN*^*234R*^, an upstream fragment was PCR amplified with primers MP67 and KN_K23R_rev and a downstream fragment with primers KN_K4R_for and MP68. For *pGIIB-KN:: venusYFP:KN*^*2R*^, an upstream fragment was PCR amplified with primers MP67 and KN_K2R_rev and a downstream fragment with primers KN_K2R_for and MP68. Each full-length fragment was PCR amplified by combining the upstream and downstream fragments with primers MP67 and MP68. The PCR-amplified full-length fragment was cut with *Xba*I and *EcoR*I and in-frame subcloned into the pGIIB-KN::venusYFP expression cassette plasmid. Sequences and open reading frame were validated with Sanger sequencing (Eurofins Genomics Germany GmbH, Ebersberg, Germany). Primers sequences are shown in Table S2.

### Chemical treatment

Wortmannin treatment: Five-day-old seedlings were treated with 10 µM wortmannin (10 mM stock in DMSO, Sigma-aldrich) in liquid medium (1/2 MS, 0.1% MES, 1% sucrose, pH5.6) for 1 hour (Wm1) or 3 hours (Wm3) and followed by mild shaking in the plant growth room.

NEM treatment: 2 mM NEM (1 M stock in DMSO, Sigma-aldrich) was added followed by incubation for 30 minutes with mild shaking in the plant growth room.

Combined treatment: After 60 minutes of 10 µM wortmannin, 2 mM NEM was added in the presence of wortmannin and further incubated for 30 minutes with mild shaking in the plant growth room.

Estradiol induction and brefeldin A (BFA) treatment: 50 µM BFA (50 mM stock in DMSO/Ethanol, Invitrogen) was applied for 30 minutes, followed by 20 μM estradiol (20 mM stock in DMSO, Sigma-aldrich) for 3 hours and 30 minutes in the presence of BFA and subsequently wortmannin and NEM were added as described above.

For membrane staining, 1 µM SynaptoRed^™^ C2 (1 mM stock in DMSO, Sigma-aldrich) was applied 30 minutes before microscopic observation. When BFA-treated seedlings were prepared for microscopy, BFA was also applied 30 minutes before microscopic observation.

### Immunoprecipitation and immunoblot analysis

The weight of five-day-old seedlings was measured prior to freezing with liquid nitrogen. Thoroughly ground samples were suspended in 2.5 times more volume IP buffer (50 mM Tris pH7.5, 150 mM NaCl, 1 mM EDTA, 0.5% Triton X-100) supplemented with an EDTA-free protease inhibitor cocktail (Roche Diagnostics). 2 mM NEM was added in IP buffer except Figure 1, Figure S1A and Figure S2B (*only wortmannin* and *only NEM*). After removal of cell debris, cleared lysates were incubated with GFP-trap beads (Proteintech, USA) for 2 hours in a cold room with mild rotation. The beads were washed five times with washing buffer (50 mM Tris, 150 mM NaCl, 1 mM EDTA, 0.2% Triton X-100) supplemented with the EDTA-free protease inhibitor cocktail and resuspended in Laemmli buffer.

For immunoblot analysis of proteins from plants, T2 seedlings were harvested. Total proteins were extracted with lysis buffer supplemented with the EDTA-free protease inhibitor cocktail. Antibodies of anti-KN (rabbit, 1:4,000) (Lauber et al., 1997), anti-GFP (rabbit, 1:1,000, Invitrogen), anti-UBQ (mouse, 1:500, Santa Cruz), anti-K63 (rabbit, 1:500, Cell Signaling Technology), anti-K48 (rabbit, 1:500, Cell Signaling Technology) were used to detect the indicated proteins. POD-conjugated secondary mouse (1:10,000, Invitrogen) or rabbit (1:20,000, Agrisera) antibodies were used. Membranes were developed with a chemiluminescence detection system (Fusion Fx7 Imager, PEQlab, Germany).

### Confocal microscopy, image processing and image quantification

Live five-day-old seedlings were used for detecting fluorescence. Fluorescent images were acquired with a confocal laser scanning microscope (TCS-SP8, Leica), using Leica LAS X software (v.3.5.7.23225).

*Line scan*. The signal intensity of the scanned lines was measured using the Leica LAS X program (v.3.0.0.15697). For Figure 3Q, the signal intensities at the plasma membrane (PM) and in the cell division plane (CP) were quantified, and for each dividing cell, the signal intensity ratio of PM to CP was calculated by setting the CP intensity at 100.

### LC-MS/MS analysis and MS data processing

Six grams of 5-day-old GFP:KNOLLE seedlings were treated with wortmannin and subsequently NEM as described above. Immunoprecipitation was done with GFP-trap beads in the lP buffer containing 2 mM NEM, rinsed with the washing buffer without NEM and resuspended in Laemmli buffer for further analysis as also described above. Proteins were eluted from the washed beads and separated on a 4-12% gradient NUPAGE Novex Bis-Tris Gel (Invitrogen). The gel section of interest was cut for protein digestion. Double digest with Trypsin and Chymotrypsin was performed as described previously (Borchert et al 2010) with the exception that chloracetamide was used for reduction and digest was performed in the presence of 10mM CaCl_2_. Extracted peptides were desalted using C18 StageTips (Rappsilber et al 2007) and subjected to LC-MS/MS analysis. Peptide analysis was performed on an Easy-nLC 1200 system coupled to an Orbitrap QExactiveHF mass spectrometer (Thermo Fisher Scientific) as described with slight modifications (Aly et al., 2023): peptides were eluted using a 49 min segmented gradient of 10–33-50-90% HPLC solvent B (80% ACN in 0.1% formic acid). MS data was processed with MaxQuant software suite version 2.2.0.0 (Cox & Mann, 2008). Database search against *Arabidopsis thaliana* database, (Uniprot, 138653 entries, 2024) was performed using the Andromeda search engine (Cox et al, 2011), which is a module of the MaxQuant. iBAQ values (Schwanhäusser et al., 2011) were enabled, minimal peptide length for identification was set to 5 amino acids. In database search, full tryptic specificity was required and up to three missed cleavages were allowed. Carbamidomethylation of cysteine was set as fixed modification, whereas oxidation of methionine and acetylation of protein N-terminus were set as variable modifications, as well as GlyGly(K), Phospho(STY), and NEM(C). Mass tolerance for precursor ions was set to 4.5 ppm and for fragment ions to 20 pm. Peptide, protein and modification site identifications were reported at a false discovery rate (FDR) of 0.01, estimated by the target/decoy approach (Elias and Gygi, 2007). For protein group quantitation, a minimum of two quantified peptides were required.

### Intensity quantification of the immunoblots

The immunodetection images were quantified with the ImageJ Fiji program (NIH).

### Box plot

The box plots with jittered data points and the group box plots were generated using the ggplot2 package (Wickham, 2009) in the RStudio program (v.1.4.1106).

### Statistical analysis

Unpaired Student’s t-test was used for Figure3Q. All *P* values were calculated with two-tailed tests.

## Supplemental information

Supplemental Figure S1 (related to Figure 1).

Supplemental Figure S2 (related to Figure 2).

Supplemental Figure S3 (related to Figure 2).

Supplemental Figure S4 (related to Figure 2).

Supplemental Figure S5 (related to Figure 2).

Supplemental Table S1. Complementation tests.

Supplemental Table S2. Primer sequences.

## Acknowledgements

We thank Alex Obholz for contributing to the complementation tests and Drs. Martin Bayer and Farid El Kasmi for critical reading of the manuscript. This work was funded by the Deutsche Forschungsgemeinschaft (DFG, German Research Foundation) – Projektnummer 516734699 (grant JU 179/27-1 to G.J.). The confocal laser-scanning microscope (TCS-SP8, Leica) used in this study was funded by an installation grant from the Deutsche Forschungsgemeinschaft (INST 37/819-1 FUGG).

## Author contributions

M.P. and G.J. conceptualized the project and wrote the original draft of the manuscript. M.P. devised the methodology and conducted the investigation. I.D.-B. and B.M. conducted the mass spectrometry analysis. M.P., I.D.-B., B.M. and G.J. reviewed and edited the manuscript. G.J. acquired the funding, provided the resources and supervised the project.

## Declaration of interests

The authors declare no competing interests.

## Notes

### Competing Interest Statement

The authors have declared no competing interest.

