## Supplemental Information for "Degradation of cytokinesis-specific Qa-SNARE KNOLLE is regulated by context-dependent ubiquitination"

Submitted to BIORXIV 13 May 2026

**Inventory of supplemental information**

Supplemental Figure S1 (related to Figure 1)
Supplemental Figure S2 (related to Figure 2)
Supplemental Figure S3 (related to Figure 2)
Supplemental Figure S4 (related to Figure 2)
Supplemental Figure S5 (related to Figure 2)

Supplemental Table S1. Complementation tests
Supplemental Table S2. Primer sequences

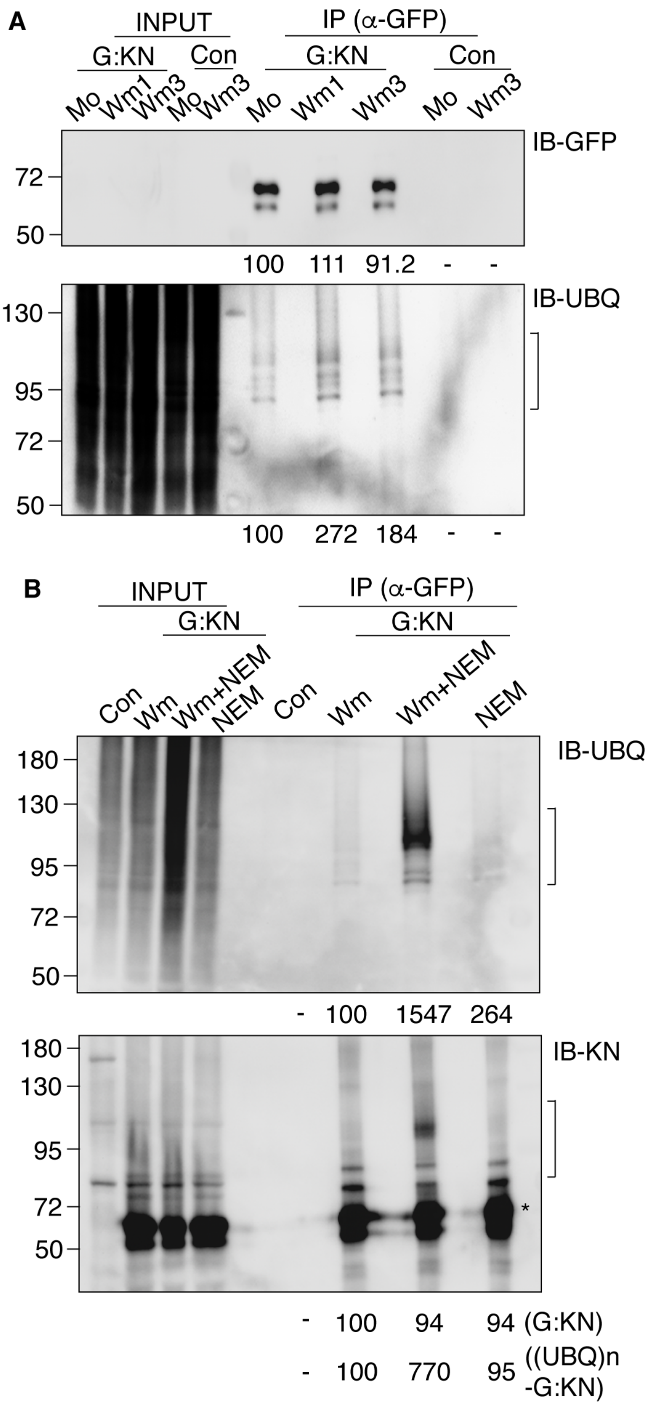


**Figure S1. Increased accumulation of ubiquitinated KNOLLE after treatment impairing endocytosis and de-ubiquitination in seedlings**

(**A**) Impairing endocytosis with wortmannin (Wm) moderately increased the level of ubiquitinated KNOLLE.

(**B**) N-ethylmaleimide (NEM) treatment affecting the activity of de-ubiquitinating enzymes had a similar effect to wortmannin on its own. The combined treatment with Wm and NEM gave a strong band also detectable with the anti-KNOLLE antiserum (double arrow; *bottom*). 'Wm+NEM' treatment included 2 mM NEM during protein extraction and immunoprecipitation. Relative signal intensity was measured by setting the signal of each control at 100 (**A**-**B**). Molecular size markers in kDa (left, **A**-**B**).

Con, non-transformed wild type; G:KN, GFP-tagged KNOLLE; Wm1, wortmannin treatment for 1 hr; Wm3 for 3 hr; Mo, mock treatment; INPUT, total protein extract; IP(α-GFP), immunoprecipitation with GFP-trap beads.

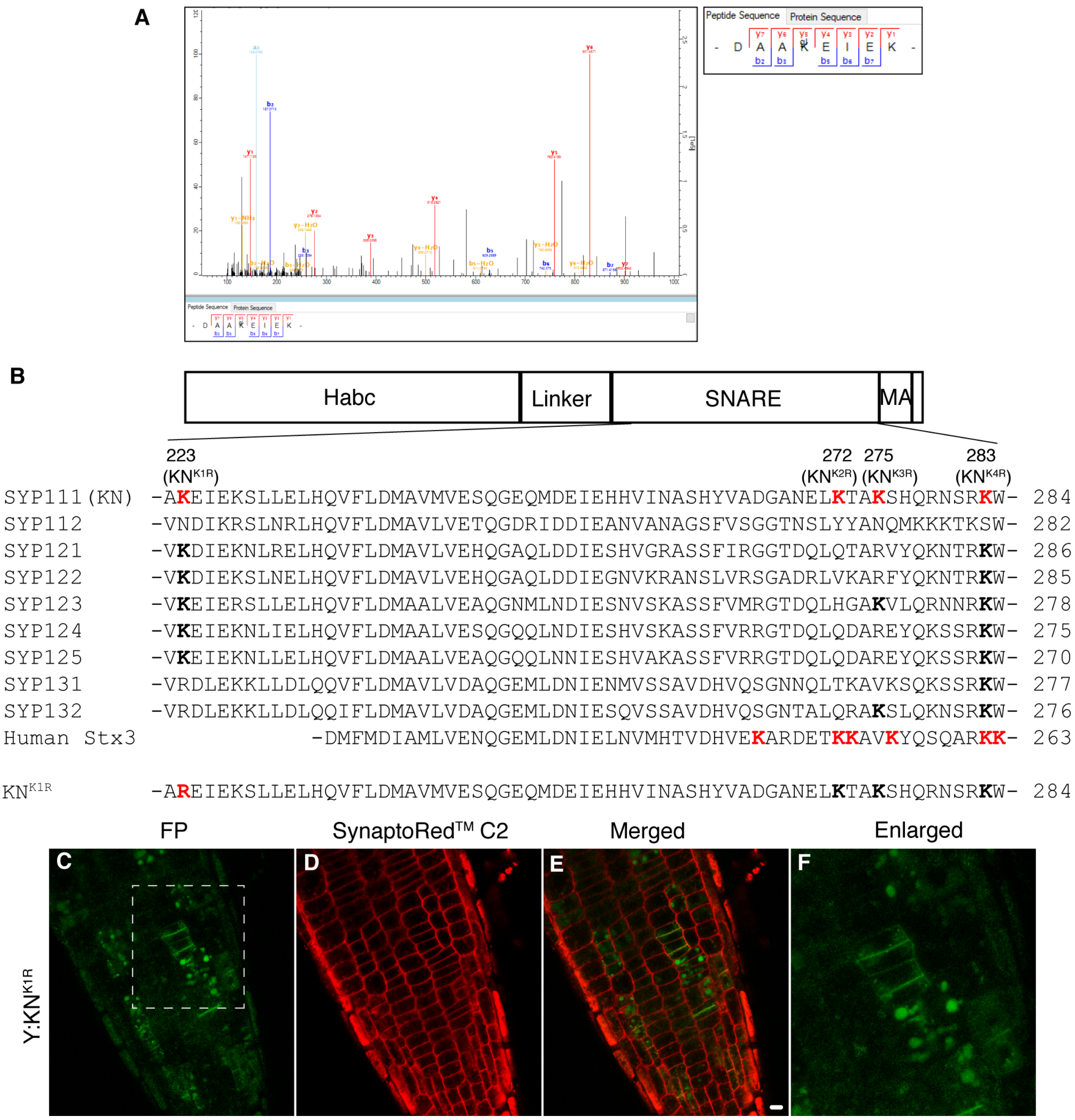


**Figure S2. Identification of four potentially ubiquitinated lysine residues in KNOLLE protein**

(**A**) IP-MS analysis of immunoprecipitated GFP:KNOLLE. *Right:* relevant sequence.

(**B**) Sequence alignment of partial SNARE domain among Arabidopsis SYP1 family Qa-SNAREs and human Syntaxin Stx3. Potentially ubiquitinated lysine residues of KNOLLE and human Stx3 are highlighted in red. *Top*, diagram of SNARE proteins: Habc, alpha-helices in N-terminal half; linker, linker between helix Hc and SNARE domain; SNARE, SNARE domain; MA, C-terminal membrane anchor.

(**C-F**) Subcellular accumulation of KNOLLE with $K_{223}$ substituted by arginine (Y:KN$^{K1R}$). Scale bar, 5 $\mu$m (**E**).

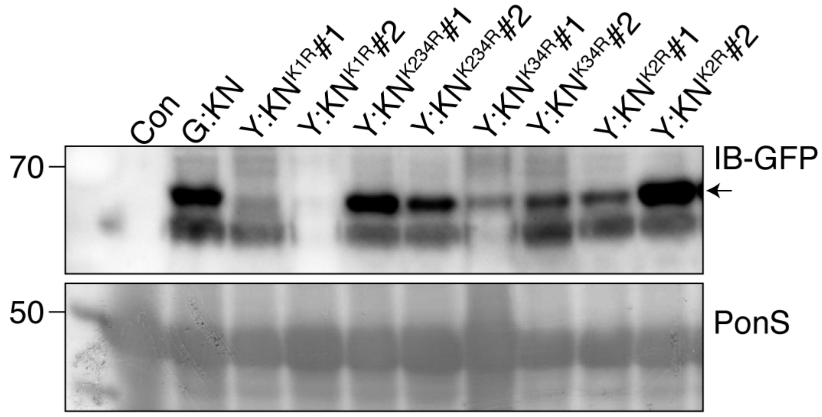


**Figure S3. Expression levels of KNOLLE in transgenic seedlings**
Protein extracts from seedlings were separated by SDS-PAGE and probed with anti-GFP antibody. Relevant bands are indicated by arrow. Con, non-transgenic wild-type control. PonS, Ponceau staining used as loading control. Molecular size markers in kDa (left).

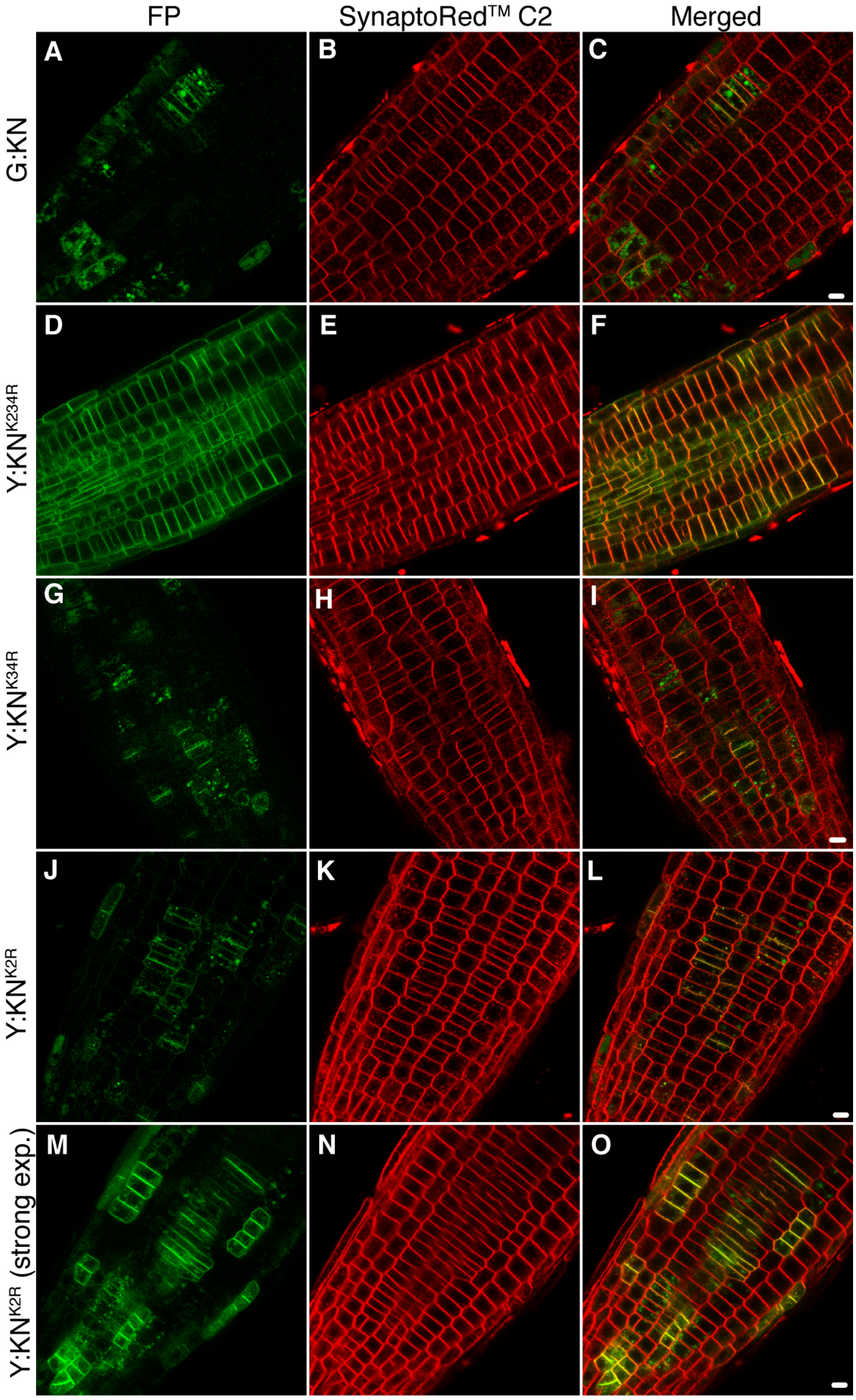


**Figure S4. Accumulation of KNOLLE variants along the seedling root (overview)**
(**A-O**) Live imaging of tagged KNOLLE variants in seedling root cells:
(**A-C**) GFP:KNOLLE wild-type protein; (**D-F**) YFP:KNOLLE with the 3 relevant lysine residues substituted by arginine (Y:KN$^{K234R}$). Note stable accumulation of Y:KN$^{K234R}$ in all root cells, like the plasma membrane marker SynaptoRed$^{TM}$ C2; (**G-I**) YFP:KNOLLE with 2 lysine residues substituted by arginine (Y:KN$^{K34R}$); (**J-O**) YFP:KNOLLE with 1 lysine residue substituted by arginine (Y:KN$^{K2R}$). Note the difference in subcellular localisation of Y:KN$^{K2R}$ between normal-expression (**J-L**) and strong-expression (**M-O**) lines.
Scale bars, 5 μm (**C**, **F**, **I**, **L**, **O**).

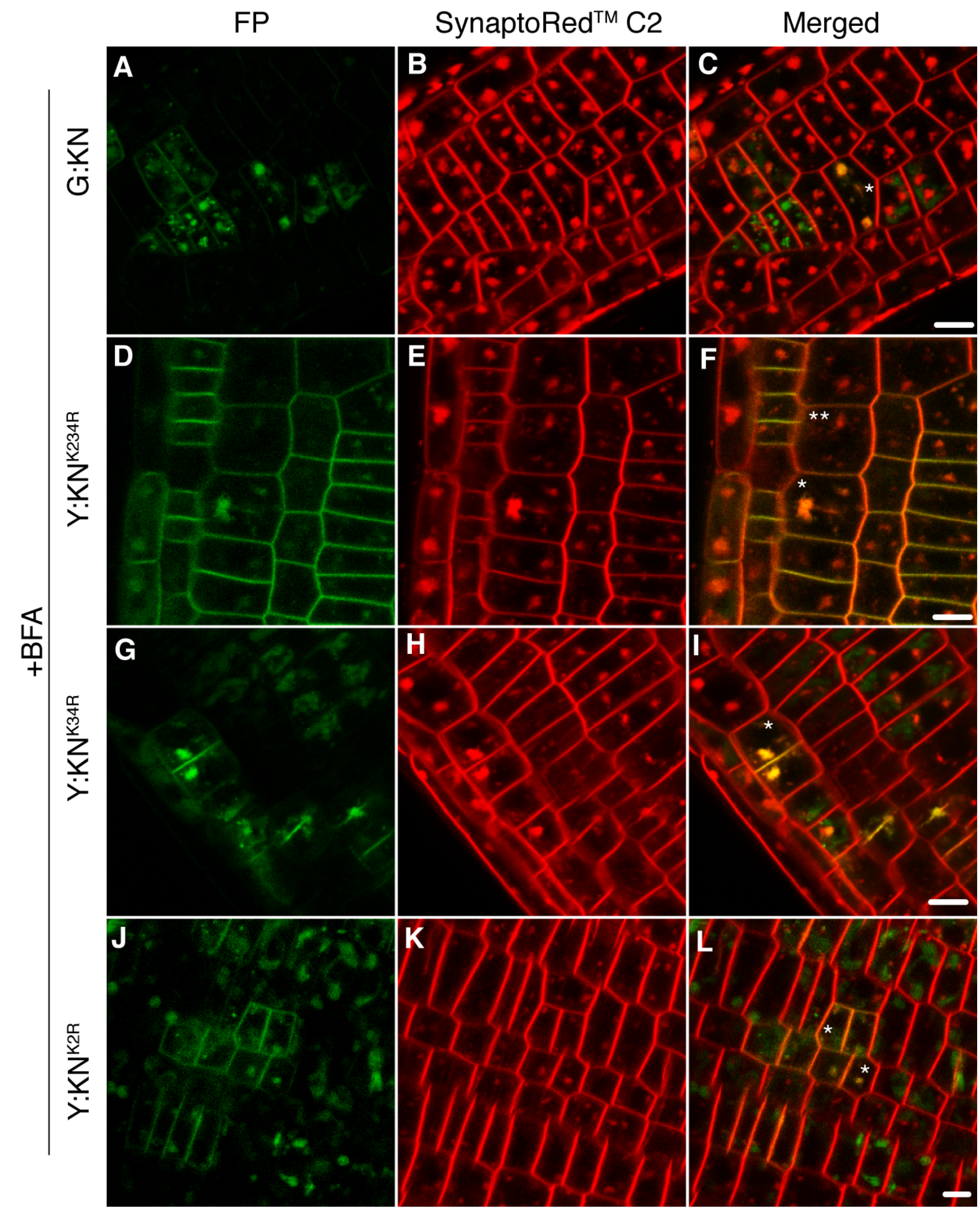


**Figure S5. Accumulation of KNOLLE substitution variants in endosomal BFA compartments**

(**A-L**) Live imaging of tagged KNOLLE variants in BFA-treated seedling root cells: (**A-C**) GFP:KNOLLE wild-type protein; (**D-F**) Y:KN$^{K234R}$.; (**G-I**) Y:KN$^{K34R}$; (**J-L**) Y:KN$^{K2R}$. Note large BFA body accumulating Y:KN$^{K234R}$ in dividing cell (asterisk in ***F***) and strongly reduced or absent YFP signal in the SynaptoRed$^{TM}$ C2-stained BFA compartments in non-dividing cells (double asterisks in ***F***). Asterisks, dividing cells (C, F, I, L); scale bars, 5 µm (**C**, **F**, **I**, **L**, **O**).

Table S1. Complementation tests

| Transgenes | T1 plants | T2 seedlings | | |
|---|---|---|---|---|
| | allele | PPTres (%)* | Phenotypic seg. (%)** | n |
| - | *kn/+* | - | 25 | - |
| *pKN::YFP:KN*$^{K1R}$*#1* | *kn/+* | 75 | 4 | 201 |
| *pKN::YFP:KN*$^{K1R}$*#2* | *kn/+* | 72 | 5 | 245 |
| *pKN::YFP:KN*$^{K234R}$*#1* | *kn/+* | 79 | 2 | 281 |
| *pKN::YFP:KN*$^{K234R}$*#2* | *kn/+* | 75 | 8 | 155 |
| *pKN::YFP:KN*$^{K34R}$*#2* | *kn/+* | 68 | 4 | 192 |
| *pKN::YFP:KN*$^{K34R}$*#5* | *kn/+* | 78 | 7 | 208 |
| *pKN::YFP:KN*$^{K2R}$*#1* | *kn/+* | 74 | 5 | 131 |
| *pKN::YFP:KN*$^{K2R}$*#3* | *kn/+* | 67 | 6 | 140 |

* The ideal segregation percentage of PPT resistance is 75% if a number of the T-DNA is one copy.

** The ideal phenotypic segregation ratio of *kn*$^{X37-2}$ mutant is 6.25% if a transgene rescues and 25% if not.

Table S2. Primer sequences

| Primers name | Sequences (5‘-3‘) |
|---|---|
| KN_XbaI_for | AAAAAATCTAGAATGAACGACTTGATGACG |
| KN_EcoRI_rev | TTTTTTGAATTCTCAAGAAGAGCTGAAACTGGT |
| KN_K1R_for | TGATGCAGCACGTGAGATTGA |
| KN_K1R_rev | TCAATCTCACGTGCTGCATCA |
| KN_K3R_rev | CTGCTGTTTCTCTGATGACTACGTGCAGTCTTCAGCTCATT |
| KN_K23R_rev | CTGCTGTTTCTCTGATGACTACGTGCAGTACGCAGC |
| KN_K4R_for | TCATCAGAGAAACAGCAGACGTTGGATGTG |
| KN_K2R_for | CTAATGAGCTGCGTACTGCAAAG*A* |
| KN_K2R_rev | TCTTTGCAGTACGCAGCTCATTAG |
| vYFP640for | ACAACCACTACCTGAGCTACC |
| x37-2 CIIIext | GGAGATTAAAAGGCTCTCTGGGACTCCGG |
| x37-2 DIIIext | ACGATGCTTGGGATGGATATGGTGGTGC |
| x37-2 DEL | CTTGATATGGCTGTGATGGTTGAAT |
